# Time is of the essence: effectiveness of dairy farm control of H5N1 is limited by fast spread

**DOI:** 10.1101/2024.11.15.623774

**Authors:** Michael E. DeWitt, Brinkley Raynor Bellotti, Nicholas Kortessis

## Abstract

The emergence and growth of highly pathogenic avian influenza(HPAI) A(H5N1) in dairy cows poses a growing threat for both the food supply and onward zoonosis to humans. Despite ongoing surveillance and control measures recommended by animal and public health authorities to limit viral spread, new herds continue to report infections. We show here that the continued spread between farms could be explained by the rapid pace of pathogen spread reported within farms, which greatly limits the potential effectiveness of these recommendations. Under reasonable surveillance strategies, we show that the time farms have to mobilize interventions is extremely limited, as few as a couple days and typically less than a week. Our findings suggest that passive surveillance measures, such as detection of H5N1 via weekly bulk milk testing, comes too late such that most infections have already occurred. For current interventions to be valuable, more sensitive and extensive surveillance is needed and an emphasis should be placed on biosecurity practices rather than reactive practices.

## 1 Introduction

Recent expansion of highly pathogenic avian influenza(HPAI) A(H5N1) to the Americas and in various, new mammalian host species is a worrying development in the history of the pathogen [1]. Perhaps the most striking development is H5N1’s emergence in dairy cows, an event that threatens a crucial component of the US food supply and opens a new route of zoonotic spillover risk to humans[2], especially given the frequent and intense contact between this new host and humans. H5N1 was first documented in dairy cattle on March 25, 2024 in Texas [3], but phylogenetic evidence suggests continued spread since November 2023 [4]. As of August 26, 2024, H5N1 is documented in 192 herds in 13 states in the US [5]. As of November 8, 2024 this has growth to 473 herds in 15 states despite biosecurity recommendations from the United States Department of Agriculture (USDA)(Figure 1). Identified cases in humans so far have been mild [6, 7], but prior H5N1 outbreaks in humans caused severe disease with high case fatality rates [8, 9]. Infected dairy cows, however, show marked decreases in milk production [3, 10, 11]. To limit economic losses and risk for spillover to humans, transmission must be limited within and between farms. The United States Department of Agriculture (USDA) thus recommends a number of biosecurity and control measures to this end [12]. However, spread has continued with few indications these actions are sufficient to quell H5N1 spread in dairies. Here we show that the rapid spread of H5N1 within farms suggests that these measures, such as batch milk testing methods enacted by the Colorado Department of Agriculture (CODA) [13], typically come too late to be effective.

**Figure 1:**
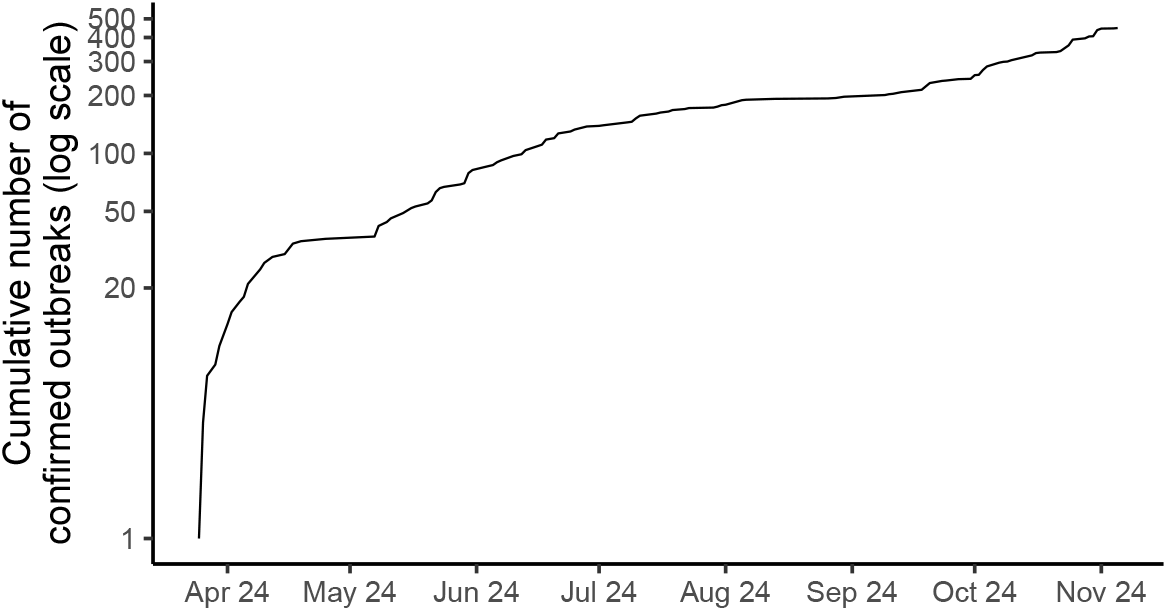
Epidemic curve of H5N1 outbreaks at the farm level. The approximate linear relationship between cumulative infections and time on the log scale suggests weakly limited, exponential growth across farms in the US. Outbreak numbers current as of November 8, 2024. The initial, rapid rise in farm outbreaks early in the epidemic is suggestive of increased detectability.

The risk of (re-)emerging diseases is often evaluated with the basic reproduction number, *R*_0_, the average number of secondary infections caused by an infection in a fully susceptible population. *R*_0_ informs the probability of pathogen establishment [14] and the effectiveness of control strategies [15]. Early, conservative estimates of H5N1 *R*_0_ on farms (between 1.2 and 1.5 [16]) suggest spread that is controllable, similar to rabies, Ebola, and Marburg viruses; pathogens that cause serious disease, but are not at high risk of uncontrollable spread. However, continued spread of H5N1 among dairy cows, despite interventions, suggests other factors are at play.

One hypothesis for the continued spread between and within farms is that mildly symptomatic, pre-symptomatic, or/and asymptomatic transmission between and within farms may be occurring. The proportion of transmission that occurs before symptoms or clinical signs appear has been directly tied to the controllability of an outbreak [17]. When infectiousness begins prior to the presentation of symptoms or other clinical signs syndromic surveillance will have a delay in detecting the outbreak. Similarly, if cows do not show any signs of illness, transmission may go on within a farm unnoticed until symptoms appear. Experimental infection studies of lactating dairy cows have shown that infected lactating cows immediately begin to have increased body temperature, reduced feeding, and reduced milk production within the first day after inoculation [10, 11]. From these infection studies we cannot rule out asymptomatic transmission occurring from non-lactating animals nor can we infer how well these studies recapitulate infections on farms. For example, Halwe and colleagues were able to detect nasal viral shedding among calves [11] which could contribute to transmission dynamics on a single farm. However, Bellotti et al considered a Susceptible, Exposed, Infected, Recovered framework when examining published outbreak reports and found that the incubation period should be on the order of several hours [16] when fit to the data with reasonable parameters, consistent with the infection studies. Given the existing evidence, it appears that asymptomatic and or pre-symptomatic transmission does not explain why outbreaks on farms unfold so quickly.

During outbreaks, time is a crucial resource that is not reflected by *R*_0_. Another measure complementary to *R*_0_ is epidemic speed. Epidemic speed, denoted by *r*, quantifies how fast pathogens spread [15]. While *R*_0_ provides insight into the effectiveness of longer term interventions such as vaccination targets, *r* is valuable in determining how much time any intervention has before it becomes ineffective in stymieing an outbreak. It is well known that earlier detection coupled with earlier interventions will result in fewer subsequent infections, but precisely how fast these detection and how strong the intervention must be is critically important when designing both the surveillance strategy and the intervention. Current estimates of *r* values for H5N1 in cattle (between 0.25-0.45 infections/day) are similar to pathogens that are difficult to control, including SARS-CoV-2, H1N1, Varicella, and Mumps (Fig. 2A). Isolation procedures, which are currently recommended by USDA for dairy farms, will prove ineffective if transmission occurs before infected cows can be identified and isolated. Outbreaks can occur for some time before being detected, which limits the time for implementation of a control (Fig. 2B). Thus another hypothesis for why the current outbreaks of H5N1 on dairy farms is that the speed of the outbreaks are such that the outbreak is nearly over by the time we detect the outbreak.

**Figure 2:**
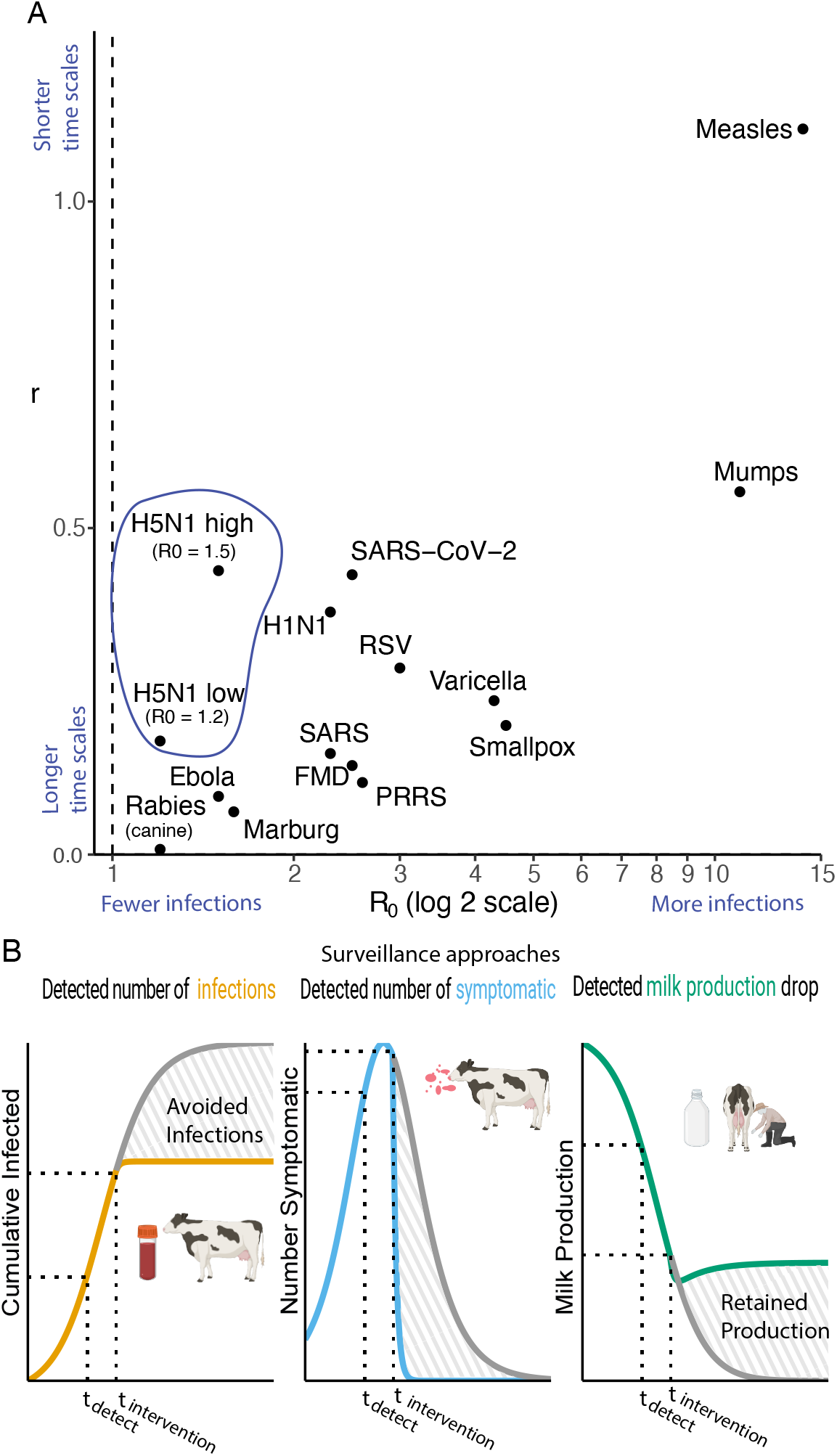
Surveillance and control strategies require knowledge of both the basic reproduction number and the intrinsic growth rate. (A) A plot comparing zoonotic and human pathogens by the basic reproduction number, *R*_0_, and the speed of an outbreak, r. (B) An overview of the three surveillance strategies examined. The outbreak is detected at some time *t*_detection_ when the detection threshold is met. An intervention identifying and isolating newly infectious cows is employed with some delay at time *t*_implementation_. The shaded regions represent either the infections avoided or the retained milk production while the gray line represents the value in the absence of intervention. Created with Biorender.com

To explore whether the pace of spread of H5N1 on dairy farms is limiting control of the disease, we have constructed a deterministic compartmental model to examine the impact of different surveillance and intervention strategies on the number of infections avoided. We assume a best case scenario that once H5N1 is detected on a farm, a highly stringent control and isolation policy is implemented that completely reduces further spread. We considered three surveillance approaches varying in sensitivity of detection and cost: monitoring cows for infection using sensitive, early detection methods (e.g., molecular testing), monitoring cattle for clinical signs (symptomatic), and monitoring changes in herd milk production. For each surveillance approach, we consider different detection thresholds, which roughly equate to the inverse of sampling intensity (e.g., detecting H5N1 when 5 cows are infected requires greater surveillance coverage than when 50 cows are infected). We aim to 1) characterize how effective any intervention can be as a function of the time between detection and mobilization of the intervention, and 2) determine how quickly a farm has to deploy a given intervention before it will fail to contain the outbreak.

## 2 Methods

### Modeling infection dynamics

We use a modified Susceptible-Infected-Recovered ODE model to represent the infectious disease dynamics of H5N1 on a single dairy farm with *N* cattle. Cows are organized into four classes representing the infectious disease dynamics as well as the manifestation of clinical symptoms. Susceptible cows are represented by class *S*. Susceptible cows become infectious (class *I*) with force of infection *βI/N*. Infectious cows no longer are infectious after a time 1*/γ* and join class *B*, in which cows are no longer infectious but still exhibit clinical symptoms. After a duration 1*/κ*, cows recover fully (i.e., are not infectious nor symptomatic) and join class *R* (Eq. 2 - 4). The total herd size is *N* = *S* + *I* + *B* + *R*. As discussed above, there is limited evidence supporting a latently infected class so it was ommited from the differential equations.

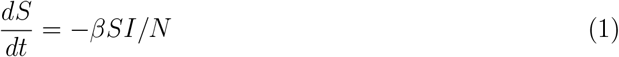

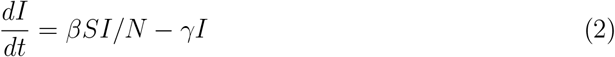

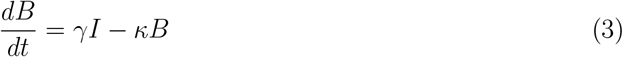

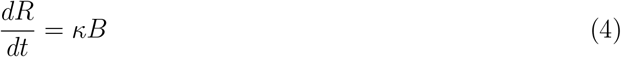

Cows are regularly phased out of milk production and replaced with cows early in their productive lifetimes (2-4 years [18]). Given the short time period of outbreaks reported on farms (often less than a month [19, 20]), we did not include population turnover because the rate of cow replacement is so small on a month scale so as to have negligible effects on our results. We also do not include disease induced mortality because mortality is relatively rare and when it occurs, it does so long after infection [3, 12]. As such, disease induced mortality also has negligible effects on the short-term outbreak dynamics we model.

The model assumes mass action infection dynamics of a well-mixed group of hosts. Dairy cows are often housed in large groups and have dense contact with one another, especially during milking and feeding operations. Therefore, the we feel that this modeling framework appropriately captures the observed disease dynamics. The relevant parameters are *β*, the effective contact rate, *γ* the rate of loss of infectiousness (1*/γ* is the average duration of infectiousness), and *κ* is the rate of recovery following loss of infectiousness. Symptoms come in many forms, the most common being a drop in milk production, but the total duration of symptoms is *γ*^−1^ + *κ*^−1^.

In this model, the basic reproductive number is *R*_0_ = *β/γ* and the rate of spread of infectious individuals is *r* = *β ™ γ* [15]. We fit parameter values in this model according to methods described by Bellotti et al.[16] and shown in Table 1. These parameters capture the basic qualitative features of observed disease course from published outbreak reports of H5N1 among dairy cattle [19, 20]. Data on spread within farms are sparse, but data on the duration of an outbreak, the fraction of cows infected, the duration of symptoms, and the timing of peak infection are available. We used these data to fit *R*_0_ and *γ* and then calculated *β* as

**Table 1:** Parameters used to simulate an outbreak of H5N1 on a single farm in dairy cattle.

| Parameter | Value | Source |
| --- | --- | --- |
| $R_0$ | 1.2 | [16] |
| $\gamma$ | 0.87 days<sup>-1</sup> | [16] |
| $\kappa$ | 0.142 days<sup>-1</sup> | [16] |
| $m$ | 100 (lb/days) | Input (Baseline value). |
| $m'$ | 50 (lbs/day) | [10] |
| $m_R$ | 75 (lbs/day) | [10] |

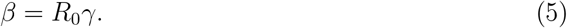

After estimating *β* and *γ*, we used reports of symptomatic duration from individual cows [10] and calculated *κ* as (symptomatic duration − *γ*^−1^)^−1^. Briefly, parameter values used in the model are consistent with an *R*_0_ of 1.2, indicating that an average cow infects 1.2 additional cows in a fully susceptible population. Note that these parameters represent the conservative lower bounds for *R*_0_. In our model fitting, *R*_0_ is primarily controlled reports of the total fraction of cattle infected. However, this reporting is subject to low ascertainment because of case reports based on symptoms, rather than true, identified infections. More-over, early reports indicate that many non-lactating cows are uninfected [10, 19], suggesting that the estimated *R*_0_ includes many cows that are not technically susceptible. Last, farms typically have pre-existing protocols to isolate suspected infected cows for other pathogens and introduce enhanced cleaning protocols (e.g., bovine mastitis [21]) which could bias existing outbreak reports downward as some interventions were employed to reduce the spread of infection. We believe that our estimate, while biased, is nonetheless conservative for the question of intervention speed on the farm because including such demographic differences among cows increases the *R*_0_ and *r* values above that used here. With faster rates of spread, control is more difficult.

### Impact of infection on milk production

Existing studies published from outbreaks[3, 19, 20, 22] and infection studies [10, 11] report reduced milk production during infection and recovery. These studies find that milk production of infected cows typically fails to return to the pre-infection baseline. To capture these dynamics, we assume a baseline milk production, *m*, of 100 lbs d^−1^. We then assume that milk production declines to some value, *m*^*′*^ while in classes *I* and *B*. Ultimately, milk production among recovered cows then recovers to some new value *m*_R_, where *m > m*_R_ *> m*^*′*^. We track milk with the compartment *M*, which quantifies cumulative milk production over time. Milk production at any time is

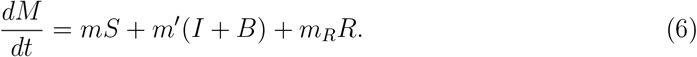

### Surveillance approaches

We model three different surveillance approaches by assuming that H5N1 is detected on a farm by monitoring different compartments in our model (Eqs 2-4,6). For each monitoring compartment, we define a “detection threshold”, which is an assumed level of change, caused by the pathogen, in a compartment that is sufficient to allow for reliable detection of the pathogen. Larger detection thresholds reflect less sensitive surveillance methods. For each surveillance method and detection threshold, we simulate an initial outbreak and measure *t*_detection_, the time since the initial infection at which the detection threshold is reached. Different detection thresholds and surveillance strategies thus map to different times during the outbreak, which influences the effectiveness of any intervention.

We make the conservative assumption that surveillance approaches are in place before the pathogen arrives on a farm (e.g., no additional delay in setting up a surveillance program). The time to detection that we measure is thus a lower bound on the actual time to detection, which may be greater because a given farm does not have surveillance structures in place at the time of infection. Hence, any delays in surveillance only add to the delay between the time of pathogen arrival and the time of deployment of an intervention. These extra delays will only reinforce our main hypothesis that the pathogen too quickly for most reactive interventions to control H5N1 on dairy farms.

The three surveillance methods are as follows:

#### Cumulative number of infections

Detection thresholds based on cumulative infections reflects the ability of a farm to identify infected individuals, either while they are currently infectious (e.g., through PCR assays) or after they have recovered (e.g., using serologic studies). This testing approach captures individuals in the *I, B*, and *R* classes and represents the most sensitive form of detection, requiring more active surveillance approaches. This approach could be considered at the level of an individual cow or through a surrogate such as testing milk for the presence of viral RNA. Infection studies have shown that viral titers rise quickly in the milk of lactating dairy cows [10, 11].

One way to view the effect of detection thresholds is to evaluate the number of hosts that must be randomly sampled to give a certain probability that the pathogen is in the random sample. For example, if few cows have been infected, they are unlikely to be captured in a small random subset of cows on the farm, whereas the same small, random subset of cows is likely more likely to contain an infected cow if many cows have been infected. To illustrate this point, consider random sampling without of cows with some number of cumulative infections. The probability of detecting a currently or previously infected cow follows a hypergeometric distribution. In a herd with 500 cows where 25 have been infected, a random sample of only 33 cows is required for there to be more than 50% chance that the disease is detected in the sample where the testing instrument has perfect sensitivity and specificity (Table 2). However, if only 2 cows have been infected out of 500, at least 354 cows must be randomly sampled before there is better than 50% chance the disease is detected.

**Table 2:** Required number of cows to sample each day under different detection thresholds in a herd of 500 cows.

| Number infected | 50% Probability of Detection | 95% Probability of Detection |
| --- | --- | --- |
| 1 | 500 | 500 |
| 2 | 354 | 488 |
| 3 | 250 | 432 |
| 4 | 193 | 376 |
| 5 | 157 | 328 |
| 6 | 133 | 290 |
| 7 | 115 | 260 |
| 8 | 101 | 235 |
| 9 | 90 | 214 |
| 10 | 81 | 196 |
| 25 | 33 | 87 |
| 50 | 17 | 44 |

**Table 3:**
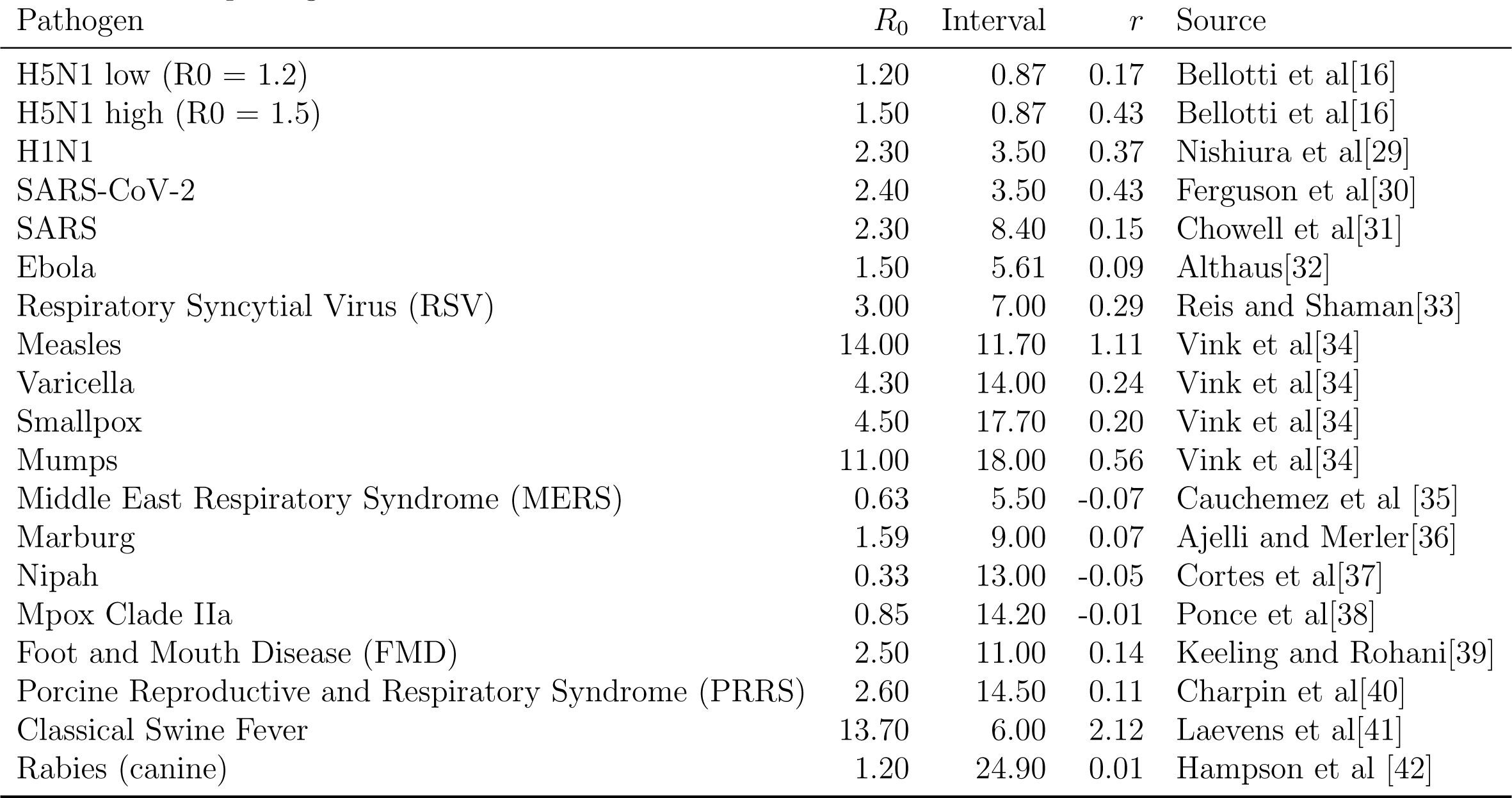
Figure 1A supporting data - Reported values of *R*_0_ and associated calculated values of *r* for selected pathogens of humans and animals.

| Pathogen | $R_0$ | Interval | $r$ | Source |
| --- | --- | --- | --- | --- |
| H5N1 low ($R_0 = 1.2$) | 1.20 | 0.87 | 0.17 | Bellotti et al[16] |
| H5N1 high ($R_0 = 1.5$) | 1.50 | 0.87 | 0.43 | Bellotti et al[16] |
| H1N1 | 2.30 | 3.50 | 0.37 | Nishiura et al[29] |
| SARS-CoV-2 | 2.40 | 3.50 | 0.43 | Ferguson et al[30] |
| SARS | 2.30 | 8.40 | 0.15 | Chowell et al[31] |
| Ebola | 1.50 | 5.61 | 0.09 | Althaus[32] |
| Respiratory Syncytial Virus (RSV) | 3.00 | 7.00 | 0.29 | Reis and Shaman[33] |
| Measles | 14.00 | 11.70 | 1.11 | Vink et al[34] |
| Varicella | 4.30 | 14.00 | 0.24 | Vink et al[34] |
| Smallpox | 4.50 | 17.70 | 0.20 | Vink et al[34] |
| Mumps | 11.00 | 18.00 | 0.56 | Vink et al[34] |
| Middle East Respiratory Syndrome (MERS) | 0.63 | 5.50 | -0.07 | Cauchemez et al [35] |
| Marburg | 1.59 | 9.00 | 0.07 | Ajelli and Merler[36] |
| Nipah | 0.33 | 13.00 | -0.05 | Cortes et al[37] |
| Mpox Clade IIa | 0.85 | 14.20 | -0.01 | Ponce et al[38] |
| Foot and Mouth Disease (FMD) | 2.50 | 11.00 | 0.14 | Keeling and Rohani[39] |
| Porcine Reproductive and Respiratory Syndrome (PRRS) | 2.60 | 14.50 | 0.11 | Charpin et al[40] |
| Classical Swine Fever | 13.70 | 6.00 | 2.12 | Laevens et al[41] |
| Rabies (canine) | 1.20 | 24.90 | 0.01 | Hampson et al [42] |

#### Instantaneous number of symptomatic cows

(i.e., cows displaying signs of infection). As part of this surveillance approach, the number of cows in the *B* compartment represent those with clinical signs that can be observed or monitored by the farm staff for signs of infection. In real-world applications, domestic livestock have some background level of infection that must be distinguishable from H5N1 infections for this to be a viable strategy. We make the conservative assumption that symptoms of H5N1 can be easily distinguished from other infections. This represents a passive form of surveillance.

Similar to the case with cumulative infections, we take detection thresholds of symptomatic cows as inversely related to the intensity of cows; a single symptomatic cow is much more likely to be sampled when there are many than when there are few symptomatic cows.

#### Milk production decrease

Representing the most passive form of surveillance, this approach examines the instantaneous milk output. The point at which the difference between some nominal baseline and the actualized output represents a threshold for detection. As in syndromic surveillance, there is some natural variation in milk production that will determine the feasibility of any detection threshold based on the well-known principles of signal to noise ratios.

We simulate a range of critical values at which point the outbreak is detected on a farm.

### Applying the intervention

In our models, interventions are modeled with an elevated removal rate of infectious cows (e.g., isolation). Let *γ*_0_ be the baseline value of *γ* and *γ*_intervention_ be the elevated rate that applies once an intervention is deployed. We assume that there is some delay, *t*_delay_, between the time of detection and the time when the intervention is mobilized. This can be summarized with the following time-dependent expression for *γ*

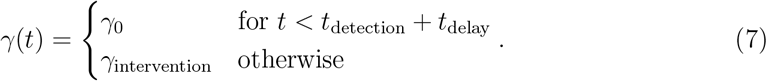

### Simulation

We simulate an outbreak over a grid of delays (*t*_delay_) and intervention intensities. We consider for identification and removal of infected cattle from 1.25 minutes (i.e., *γ*_intervention_ = 1152 per day) to 20 hours (i.e., *γ*_intervention_ ≈ 0.87 per day), the latter of which is approximately equivalent to the baseline infectious period (*γ*_0_ = 0.87 per day) and so represents effectively no intervention. For each intervention intensity, we simulated across delays ranging from 12 hours to 20 days. We then calculated the proportion of infections avoided after the implementation of the intervention as

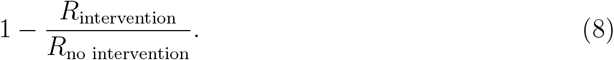

All scenarios were run with a single initial infection and 499 susceptible cows representing a herd of 500 dairy cows. We assumed *R*_0_ = 1.2, *γ* = 0.87 days^−1^, *κ* = 0.142 days^−1^, and *m* = 100 (lb/days) following Bellotti et al [16]. Following the results from Baker et al., we assumed *m*^*′*^ = 50 (lbs/day) and *m*_*R*_ = 75 (lbs/day) [10].

Code and supporting data are available at https://github.com/wf-id/h5n1speed and within a Zenodo repository [23].

## 3 Results

### Detection and deployment delays limit time to effectiveness of interventions

The most active surveillance measure (i.e., monitoring all cows, Fig. 3A) is more effective than passive forms (i.e., monitoring symptomatic cows, Fig. 3B, or milk, Fig. 3C) because H5N1 is detected earlier on farms. For example, highly sensitive monitoring for a 2% reduction in milk production is roughly equivalent to detecting the pathogen when approximately 25 cows are infected or 11 cows exhibit clinical symptoms. Furthermore, unavoidable delays between detection and intervention deployment further erode effectiveness. For example, an intervention aiming to avoid 80% of infections deployed within seven days of detection and isolates cases within 12 hours requires a detection threshold of 5 infected cows or fewer (Fig. 3A). If the same intervention could be deployed within two days, the detection threshold increases to 18 cows (Fig. 3A).

**Figure 3:**
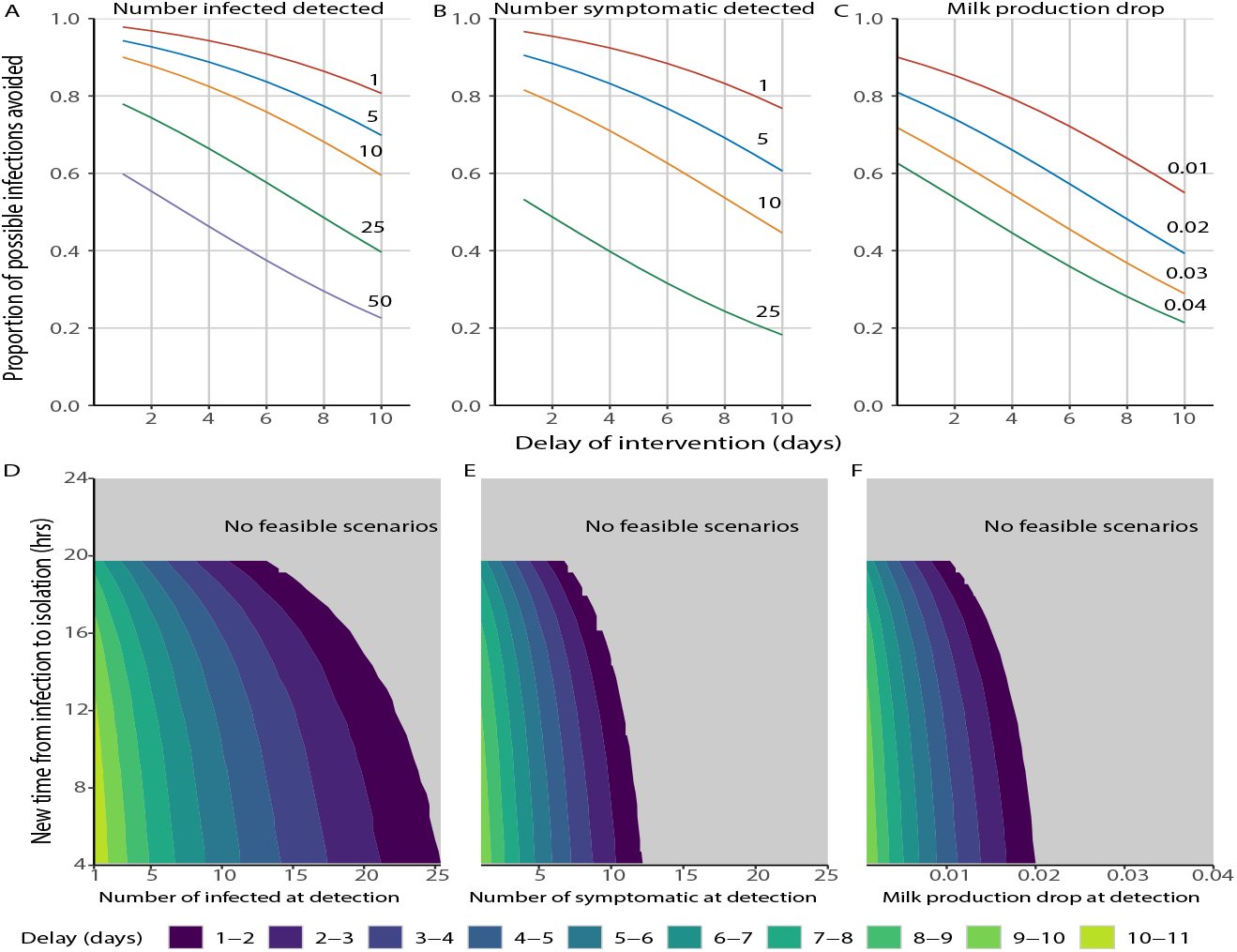
Intervention effectiveness and the role of time to effective strategy. Panels A-C reflect the proportion of the total fraction infected avoided given the strategy implementation delay and the detection threshold for an infection-based detection (A), number of clinical cows (B), and drop in milk production (C) for some intervention which can identify and isolate infections within 12 hours (*γ* = 2). Panels D-F reflect the required intensity of the intervention (y-axis, representing the effective new *γ* post-intervention) required to avoid 80% of the infections during an outbreak given the strategies’ detection threshold (x-axis) and the number of days to deploy the strategy (color contours) for number of infected cows (D), number of clinical cows (E), and milk production decrease (F).

### Passive surveillance approaches are not feasible means of control

Fig. 3D-E show how long particular intervention and detection combinations have before they lose effectiveness (here illustrated as time to avoid 80% of infections). For milk testing, an intervention must be deployed in 5 days or fewer under the highly sensitive case of detecting H5N1 when milk production declines by 0.4% (Fig. 3F). For a more conspicuous 1% drop in milk production, farms have at best 24 hours to deploy an intervention.

## 4 Discussion

Interventions should happen quickly to maximize the effectiveness and reduce the total fraction infected during an outbreak and are driven by the speed of the pathogen. Our findings suggest that more passive surveillance approaches are less effective at limiting the total number of infected during an outbreak than more active approaches. Waiting for clinical symptoms to manifest further limits the effectiveness of any intervention, a phenomena observed with asymptomatic or pre-symptomatic transmission [17].

Once an H5N1 outbreak begins on a dairy farm it is difficult to control. While *R*_0_ is far below estimates for those of Foot and Mouth Disease (FMD), we see similar dynamics with the observations from Keeling et al. where mathematical models were used to guide control strategies that treated farms as entirely infected due to the rapidity of the outbreak once a given farm had a detected case [24]. Our model suggests a similar trajectory, with control measures focused on averting an outbreak once the pathogen has been introduced on a farm being highly ineffective. The control strategy introduced by the CODA requiring weekly bulk milk testing [13] would be an ineffective detection threshold for farm-level control based our findings. Rather, our findings would suggest that more frequent bulk milk tank testing (i.e., daily tests of all milk produced) could converge on the active surveillance strategy that we explored and would allow for both prevention of infections and implementation of quarantine strategies to prevent spread to other farms.

The relationship between *r* and *R*_0_ as related through the generation interval is well described [25]. Furthermore, these relationships have been used to estimate real-time reproduction numbers in order to guide interventions [26]. However, in our analysis we approach the quantitative problem of the surveillance and rate of implementation on dairy farms. Virtually all interventions will have more impact if they is employed earlier; however how early an outbreak can feasibly be detected is a quantitative problem. Essentially, without more active surveillance approaches the outbreak is nearly over at the time of detection. This raises a larger questions about how public and animal health programs must integrate the concepts of not only asymptomatic transmission and *R*_0_, but also how speed measured by *r* can influence the surveillance strategy needed to prevent both infections within a population, but also to stymie onward infection to other populations.

The ongoing transmission of H5N1 among dairy cattle is still under intense study with many unknowns. We recognize that these epidemiologic values are likely to change as new information comes to light. However, our conclusions rely primarily on reports of the duration of outbreaks. Our simulation outbreaks are effectively over by day 40 with the peak of infections around day 19, which are consistent with epidemiologic reporting [27]. The current information, however imperfect, paints a clear picture: there is not much time to act after H5N1 can be detected.

Our work suggests that restricting pathogen introduction in the first place should be prioritized. Biosecurity practices, such as restricting the movement of animals, personnel, and other products from nearby farms, are likely to be more effective than dealing with out-breaks on farms. If these are impractical, highly sensitive detection methods are required. More frequent bulk milk testing could provide the benefits of the active surveillance strategy if the frequency is high enough and number of animals considered sufficiently large as demonstrated in Figure 3D.

While we use estimated values for *R*_0_ from prior estimates[16], these values should be taken as the lower possible bounds for H5N1. *R*_0_ taken as an average across a farm does not account for other sources of heterogeneity such as differences in susceptibility (e.g., lactating cows seem to be affected more), differences in contact structure and spatial structure (e.g., cows at different stages of lactation are in closer contact). Higher values of *R*_0_ would suggest even higher rates of speed within these groups which would further complicate possible intervention strategies to control the outbreak.

H5N1 emergence in dairy cattle represents a trial run of our pandemic preparedness. While *R*_0_ is undoubtedly important, pathogen speed should be considered when designing outbreak surveillance and intervention. For example, had H5N1 the same *R*_0_ but half the speed, farms would have approximately 17 more days to detect and implement a control strategy. At that time scale, weekly bulk testing milk would be viable.

Frieden et al. proposed the 7-1-7 framework in which an outbreak is identified within seven days, authorities notified within one day, and an intervention mobilized seven days later [28]. Such a framework takes too much time to be successful in the context of pathogens with rapid spread, such as H5N1. Pathogen speed is a critical factor determining efficacy of surveillance and intervention strategies, especially for emerging pathogens in which there is limited time to act.

## Author contributions

MD and NK designed the research; MD performed the research and data analysis. MD, BB, and NK wrote the paper. The authors declare no competing interest

## Acknowledgments

Funding was provided by the National Science Foundation (NSF Grant DMS-2327799) to NK. Computations were performed using the Wake Forest University (WFU) High Performance Computing Facility, a centrally managed computational resource available to WFU researchers including faculty, staff, students, and collaborators.

